## Supplemental Information for "Exposing the molecular heterogeneity of glycosylated biotherapeutics"

^‡^Current affiliation: Protein Analytical Development, Ascendis Pharma, CA, USA

^†^Current affiliation: Translational Pharmacometrics, Janssen, PA, USA

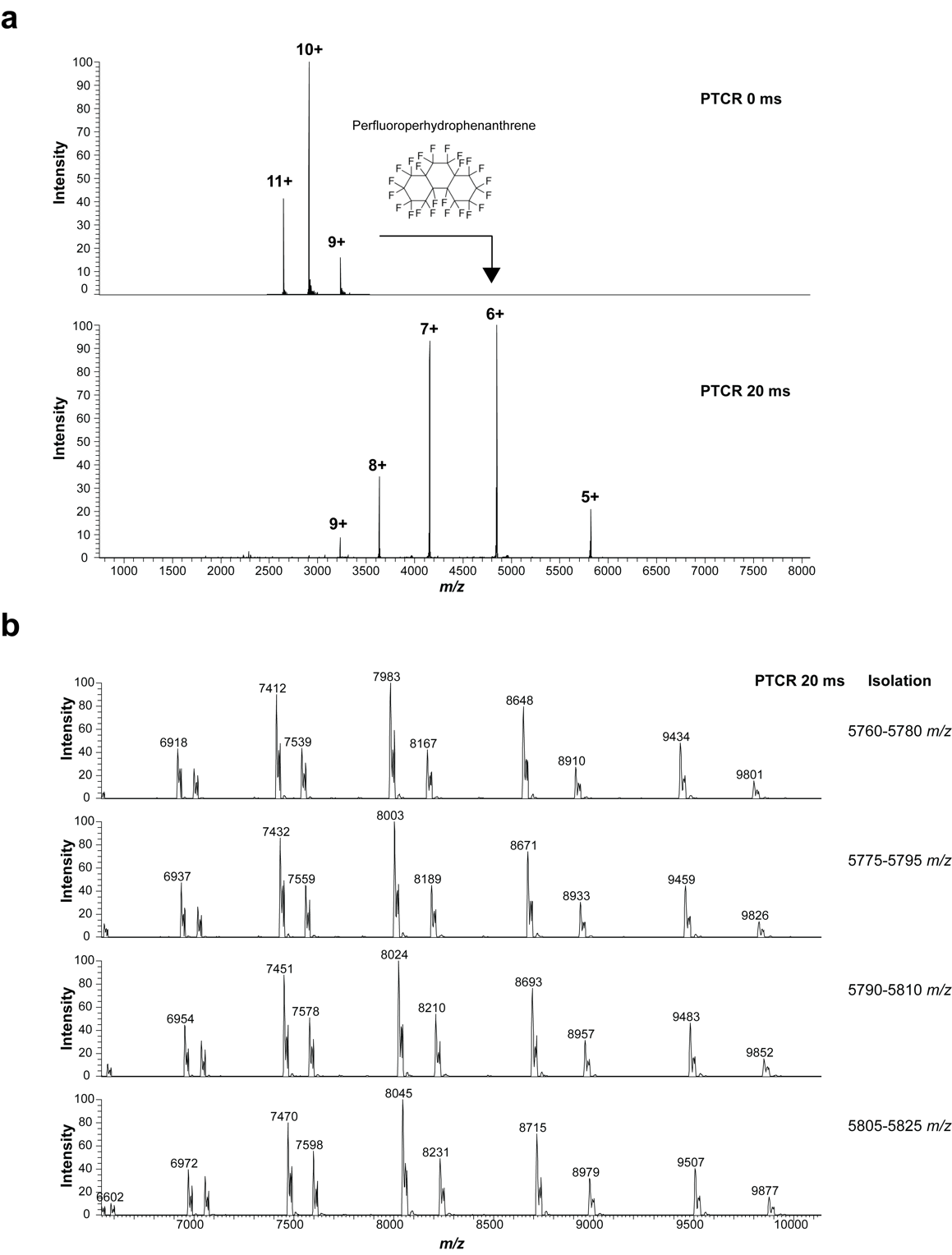

**Supplemental Figure 1.** **a**, proton transfer charge reduction, as demonstrated on bovine carbonic anhydrase II (UNIPROT: P00921), a standard protein for Mass Spectrometry method development. The reagent used is perfluoroperhydrophenanthrene (PFP). **b**, PTCR on multiple isolated ion populations of IL22-Fc, the bio-therapeutic featured in Main text **Fig. 2**, with a window of 20 *m/z* and 5 *m/z* overlap.

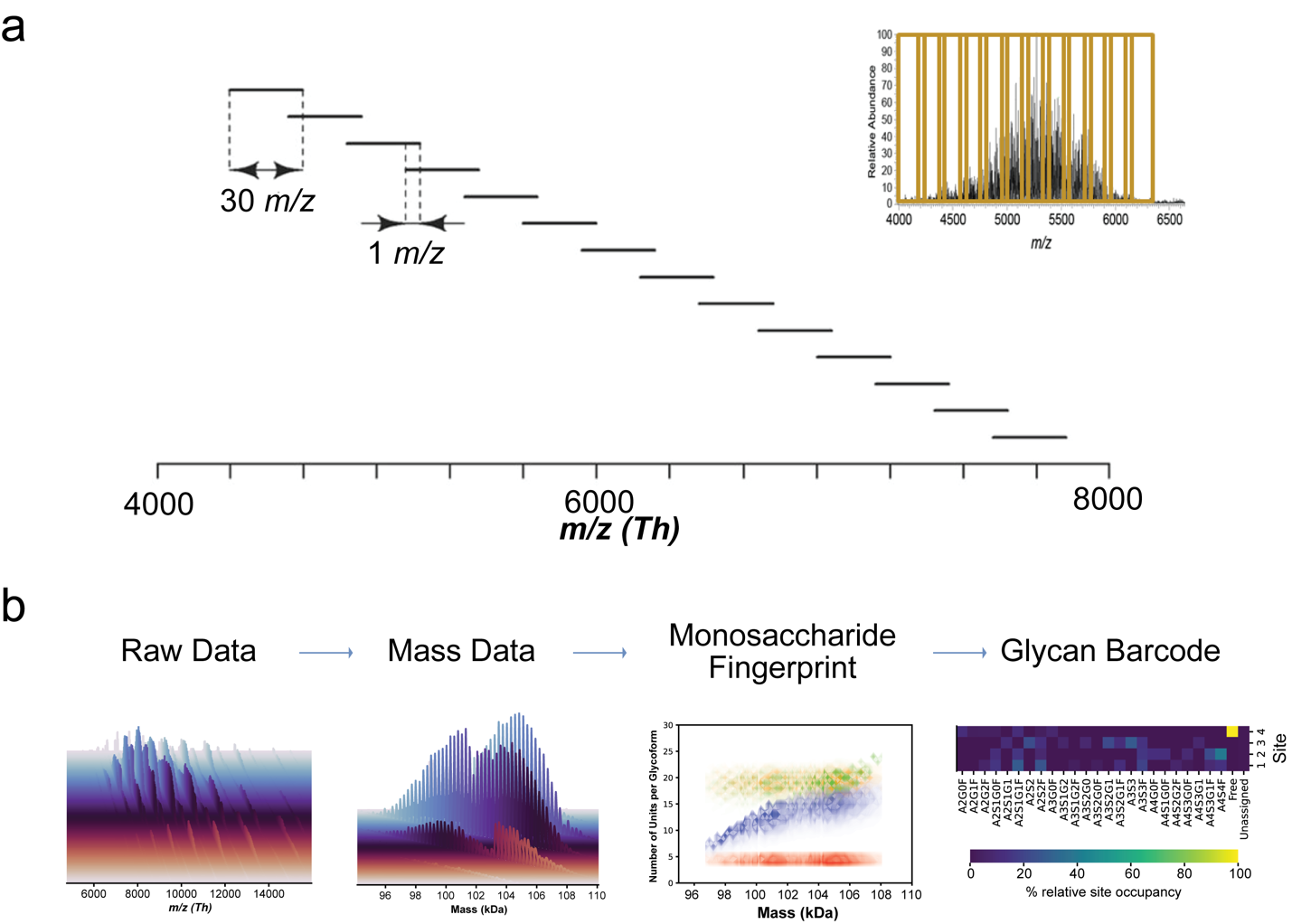

**Supplemental Figure 2.** **a**, Schematic representation of a segmented acquisition (i.e., data-independent) workflow example, with a 30 *m/z* isolation window with 1 *m/z* overlap, spanning the range of 4300-8000 m/z. The *m/z* isolation window width was chosen to be sufficiently narrow to assure that the MS^2^ PTCR spectra were relatively sparce to allow observation of the reduced charge states of the isolated glycoform ions (i.e., the glycoform-resolved spectrum). However, this was balanced with the need to minimize the total number of isolation windows stepped through in DIA-mode to acquire PTCR MS^2^ spectra which would cover the entirety (or a suitably large swath) of the *m/z* range of the glycoprotein ion signal observed in the MS1 spectrum.

**b**, General DIA-PTCR workflow for glycan fingerprinting, starting with raw data acquisition using DIA-PTCR as shown in **a**, with each colored spectrum showing the result of a single isolation event and PTCR dispersion; then the *m/z* data is processed using UniDec/UniChrom for mass deconvolution – yielding mass domain data. Intact masses for the biotherapeutic can be correlated bioinformatically with possible glycan building block structures (derived from orthogonal analyses) for ‘monosaccharide fingerprinting’ – which provides a landscape of the glycoform-resolved monosaccharide composition of the bio-therapeutic, and to generate a ‘glycan barcode’, which enables more direct comparisons of glycan composition across samples.

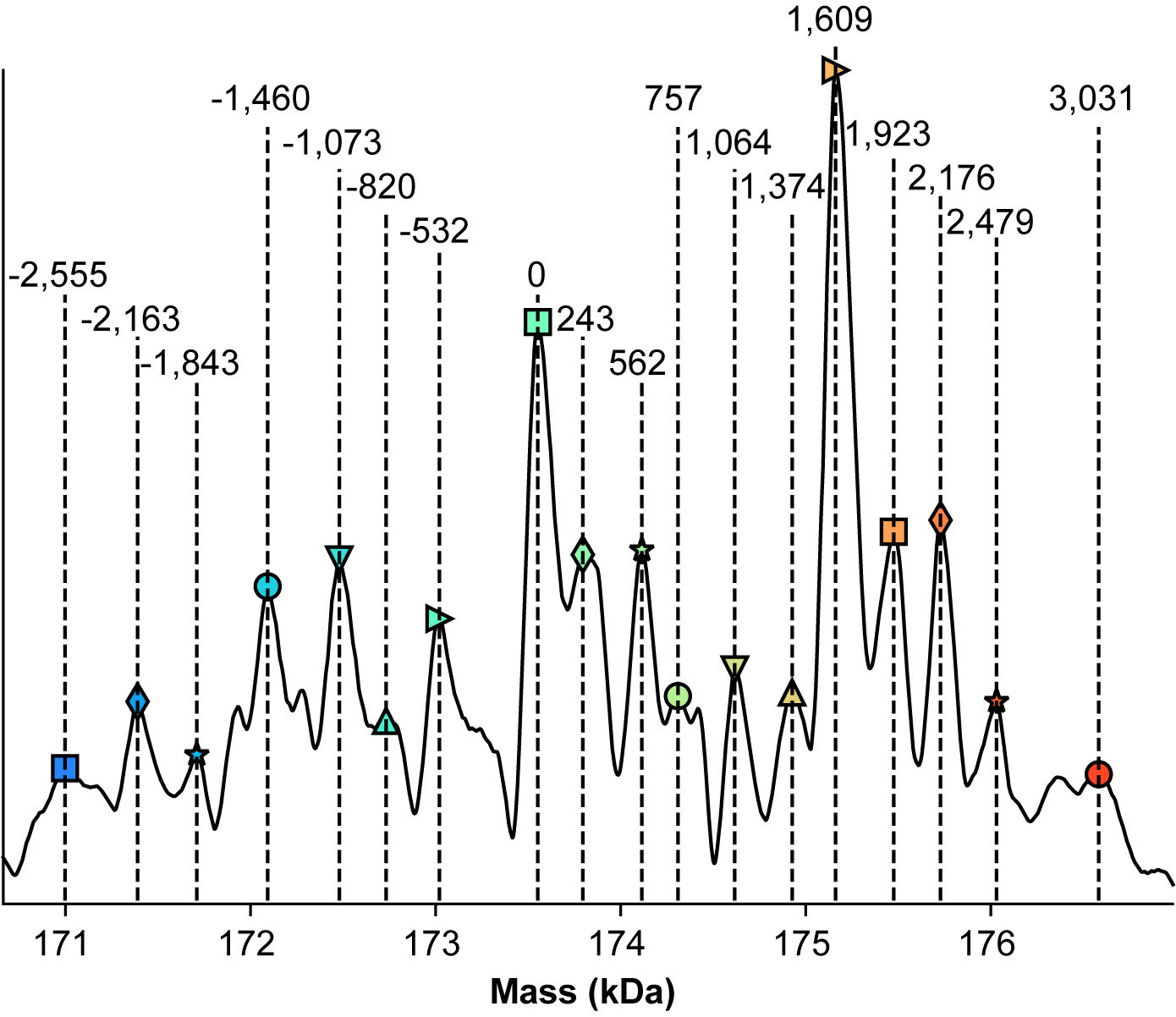

**Supplemental Figure 3.** Inset from Main Text **Figure** **1j**, showing mass differences consistent with glycosylation. Glycan annotations for mass shifts relative to the central peak in the spectrum are elucidated in the table below.

| **Mass Shift (Da)** | **Glycan** | **Theoretical Mass (Da)** |
| --- | --- | --- |
| 1,609 | G1F / dHex(1) Hex(4) HexNAc(4) | 1607 |
| 532 | Hex(2) HexNAc | 527 |
| 568 | Hex(1) HexNAc(2) | 568 |
| 243 | HexNAc(2) Hex(-1)* | 244 |
| 757 | dHex HexNAc(3) | 755 |
| 1064 | dHex(2) Hex HexNAc(3) | 1063 |
| 1374 | dHex(3) Hex(2) HexNAc(3) | 1372 |
| 1923 | dHex(2) Hex(5) HexNAc(4) | 1915 |
| 2176 | Hex(5)HexNAc(4)NeuAc(2) | 2205 |
| 2479 | Hex(6) HexNAc(6) dHex2 | 2485 |
| 3031 | Hex(9) HexNAc(7) dHex(1) | 3027 |
| 1843 | Hex(4) HexNAc(3) NeuAc(2) | 1840 |
| 1460 | G1 / Hex(4) HexNAc(4) | 1461 |
| 820 | dHex(2) Hex(2) HexNAc | 819 |

*loss of 1 Hex

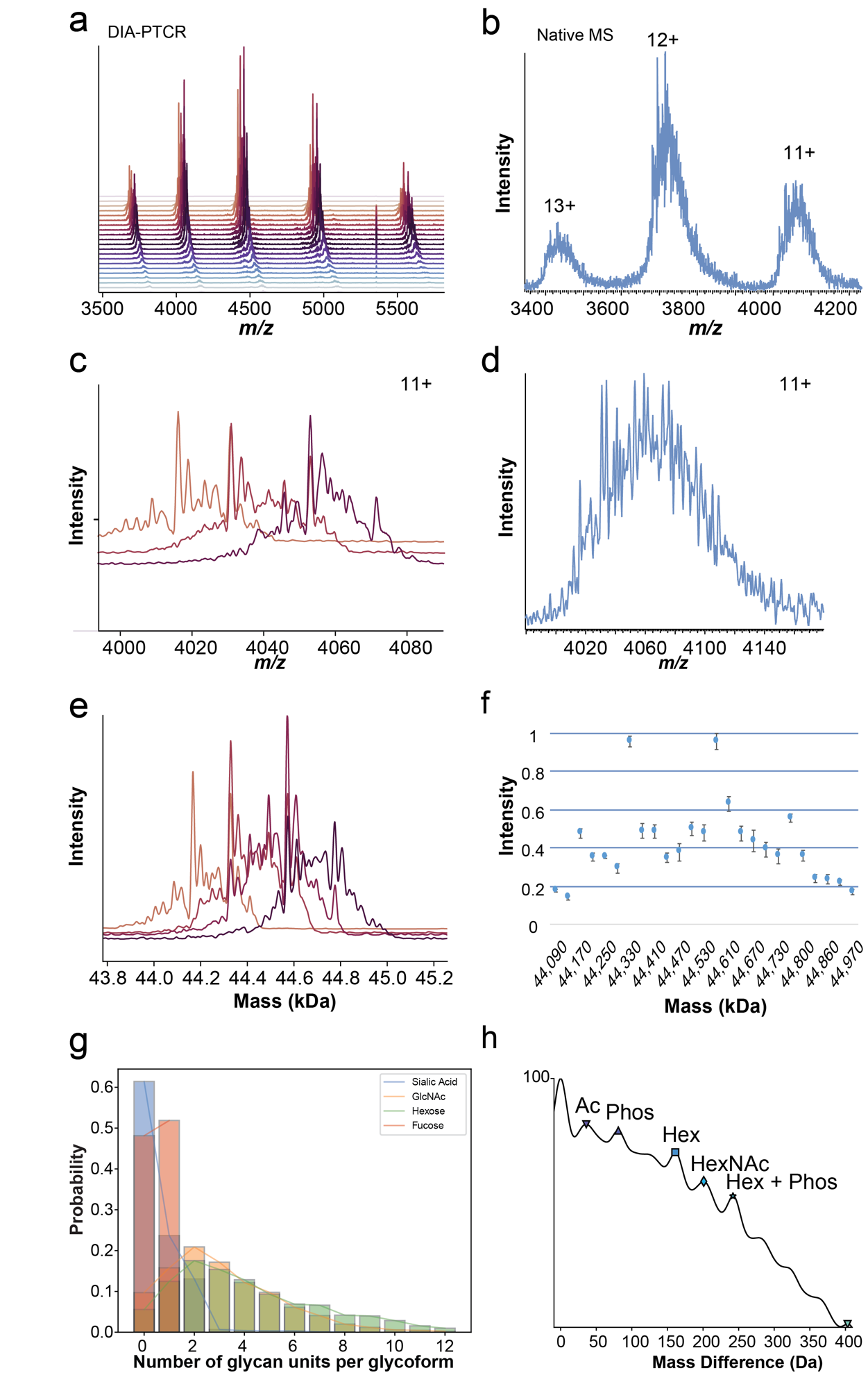

**Supplemental Figure 4**. **a**, Full stacked spectral representation of the DIA-PTCR analysis of ovalbumin from chicken egg (UNIPROT: A0A2H4Y842), acquired with a 10 Th isolation window and Orbitrap resolution of 15,000. **b**, Native MS analysis of ovalbumin without DIA-PTCR. **c**, Selected PTCR MS/MS spectra of the 11+ charge state of ovalbumin following DIA-PTCR. **d**, Zoom-in of the 11+ charge state of ovalbumin upon native MS analysis without DIA-PTCR. **e**, Selected neutral mass spectra obtained by mass deconvolution of each isolated PTCR MS/MS spectrum.  **f**, Reproducibility of DIA-PTCR analysis of ovalbumin, shown as detected masses versus relative intensity for four replicates, with error bars indicating 1 standard deviation. **g**, Number of monosaccharides per glycoforms plotted vs their probability of occurrence, determined from the DIA-PTCR dataset assuming equal probability for all matches. **h**, Autocorrelation plot generated in UniDec, showing common mass differences detected in the deconvolved mass spectrum of ovalbumin following DIA-PTCR, plotted against their probability of occurrence. These correspond to ovalbumin’s reported post-translational modifications: acetylation (Ac), phosphorylation (Phos) and glycosylation (Hex, hexose; HexNAc, N-Acetylhexosamine).

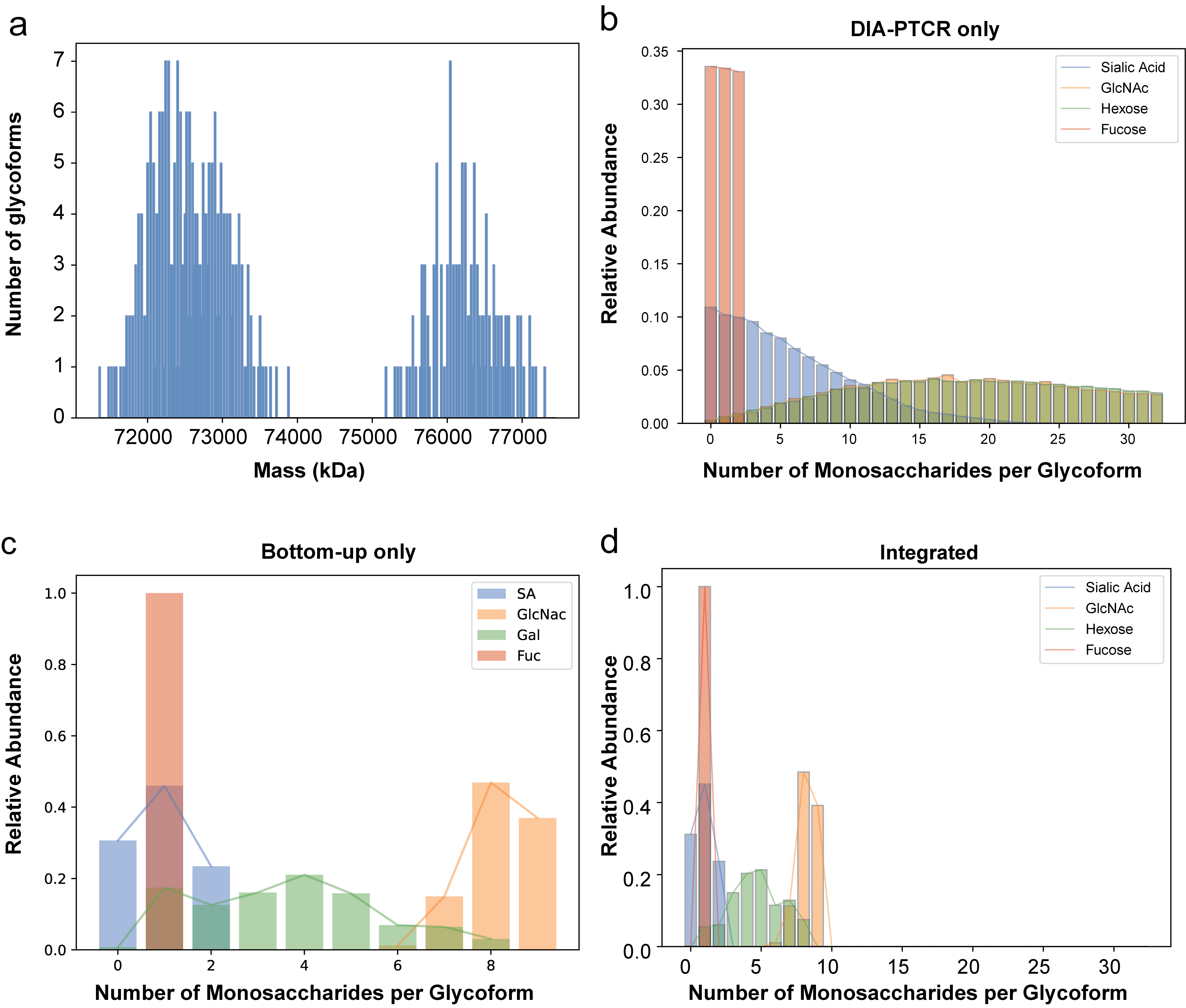

**Supplemental Figure 5.** **a**, Brute force simulation of the number of theoretical glycoform of MHCII displaying the 360 possible combinations of glycans for the four glycosylation sites on MHCII, plotted as a function of mass vs. number of glycoforms. Interestingly, the simulated bimodal distribution – due to the mixture of full-length and truncated alpha chain – is very similar to the experimental glycoform distribution detected by DIA-PTCR.

**b**, Number of monosaccharides (i.e., sialic acid, GlcNAc, Hexose and Fucose) per glycoform plotted against their relative abundance in the DIA-PTCR-only dataset. **c**, Number of monosaccharides per glycoform plotted against their relative abundance in the glycopeptide-only dataset. **d**, Number of monosaccharides per glycoform plotted against their relative abundance for the integrated DIA-PTCR and glycopeptide datasets. For more information on how these plots were generated, see Supp. Fig. 9 and the Methods section.

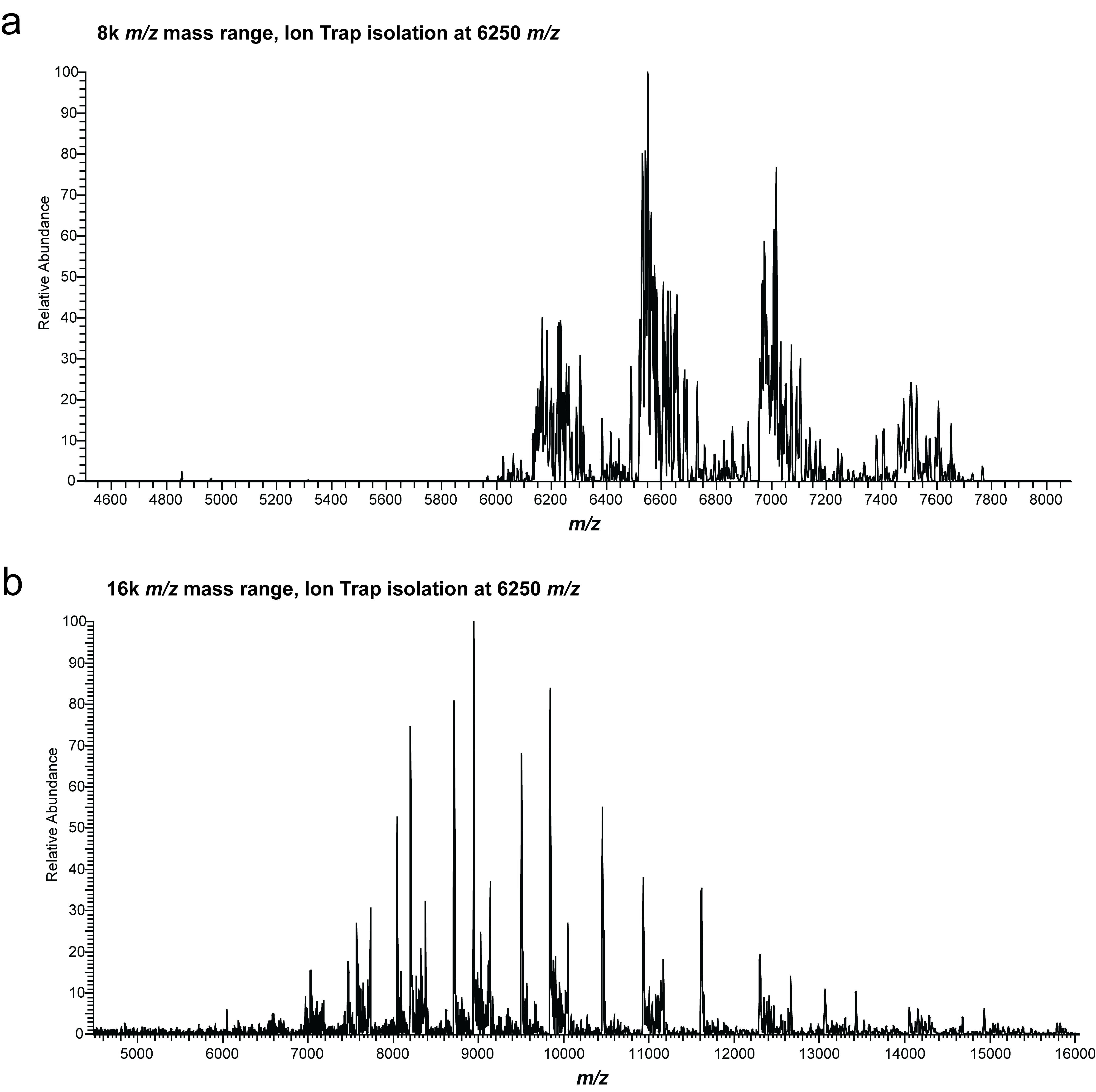

**Supplemental Figure 6.** Demonstration of PTCR of an isolated ion population at 6250 *m/z* of IL22-Fc using **a**, Orbitrap Eclipse Tribrid MS with 8000 *m/z* maximum mass range, and **b**, Orbitrap Ascend Tribrid MS with 16000 *m/z* maximum mass range. The increased mass range enables detection of lower charge states at higher *m/z* further preventing the overlap of spectral features and increasing proteoform resolution.

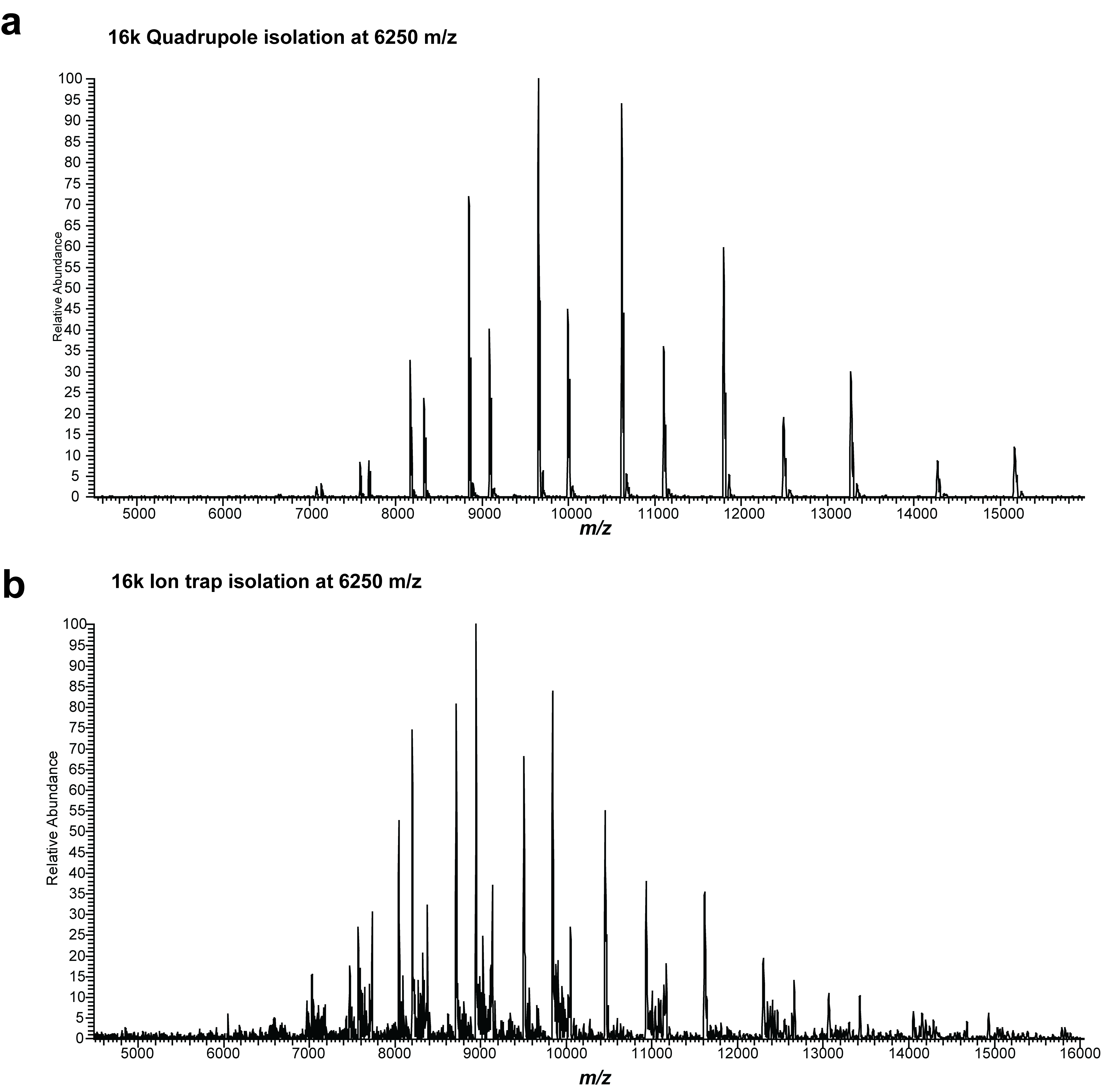

**Supplemental Figure 7.** Isolation at 6250 m/z using **a**, quadrupole with isolation width of ~20 Th or **b**, the ion trap with isolation width of 100 Th.

While both isolation methods achieve sufficient signal for glycoform identification, there are some noteworthy differences. The signal-to-noise ratio at high MS^2^ acquisition targets is significantly higher for quadrupole isolation than for the ion trap because the quadrupole is not sensitive to space-charge effects during isolation. The narrower isolation width afforded by quadrupole isolation results in fewer precursor species per isolation window, leading to fewer potential peak overlaps and overall higher signal-to-noise ratio of the isolated species due to a higher percentage of that species out of the total amount of charge isolated (similar to a boxcar enrichment).

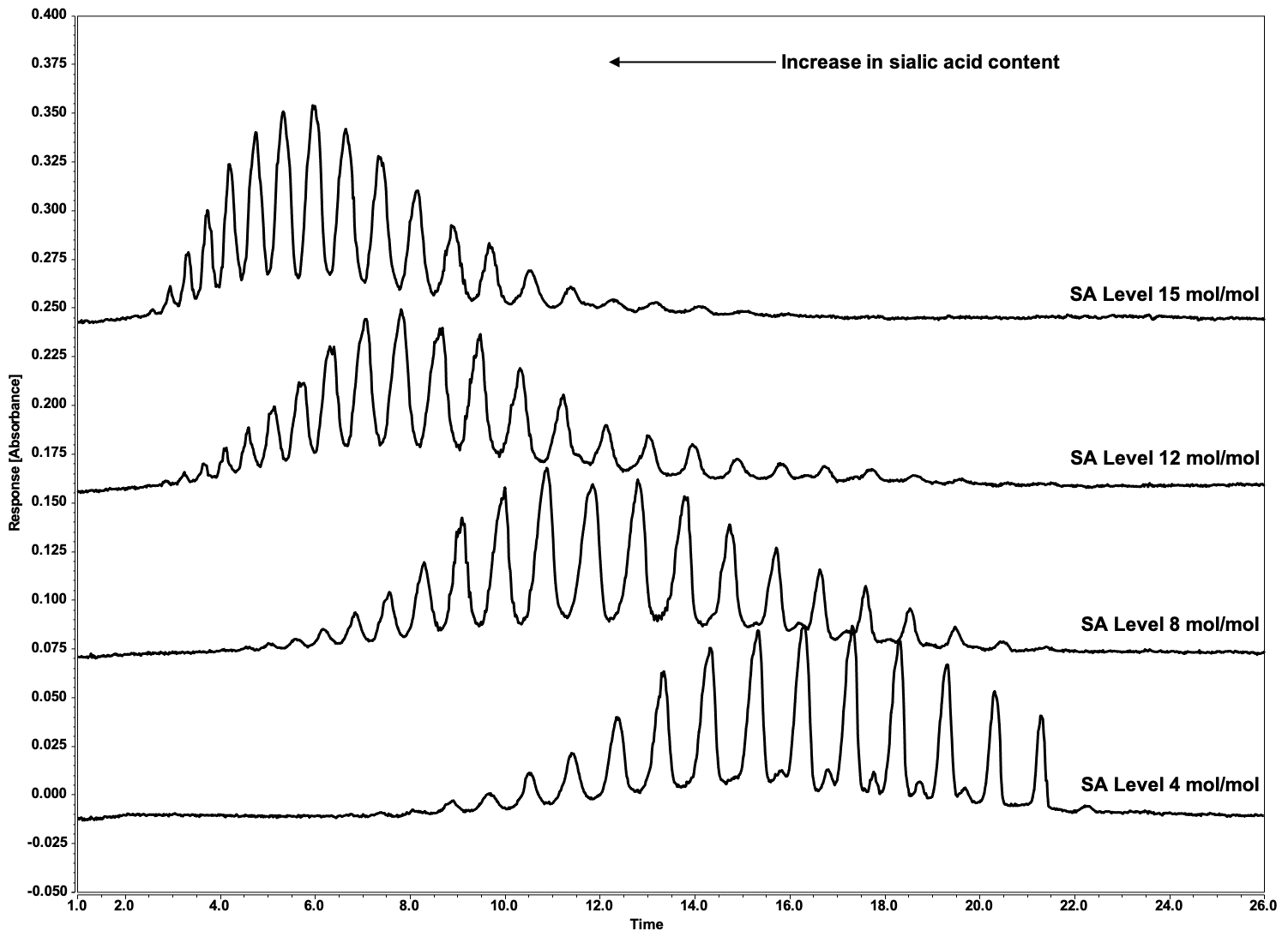

**Supplemental Figure 8.** Electropherograms of IL-22Fc Sialic Acid Variants. A purity method based on imaged capillary isoelectric focusing (ICIEF) was developed for the separation of glycoforms of IL-22Fc. The separation was found to be driven in part by the number of negatively charged sialic acids attached to each glycoform. The ICIEF glycoform charge assay was used to provide quantitative information on the distribution of sialic acid in IL-22Fc. IL-22Fc samples were treated with CpB to remove C-terminal lysine charge heterogeneity, mixed with an ampholyte solution containing urea to denature the protein, and separated in a capillary cartridge using a ProteinSimple iCE3 imaged cIEF system. The electropherograms of the IL-22Fc sialic acid variants (SA Level 4, 8, 12, and 15 mol/mol) are shown in the above figure. The negatively charged sialylated glycoforms migrate towards the acidic anolyte; thereby glycoforms with increasing numbers of sialic acids focus on the left hand side of the electropherogram. As sialic acid content increases, a shift in the glycoform distribution towards the acidic region of the electropherogram is observed indicating an increase in negatively charged sialic acid residues (as is observed by 2-AA HILIC UHPLC in next figures).

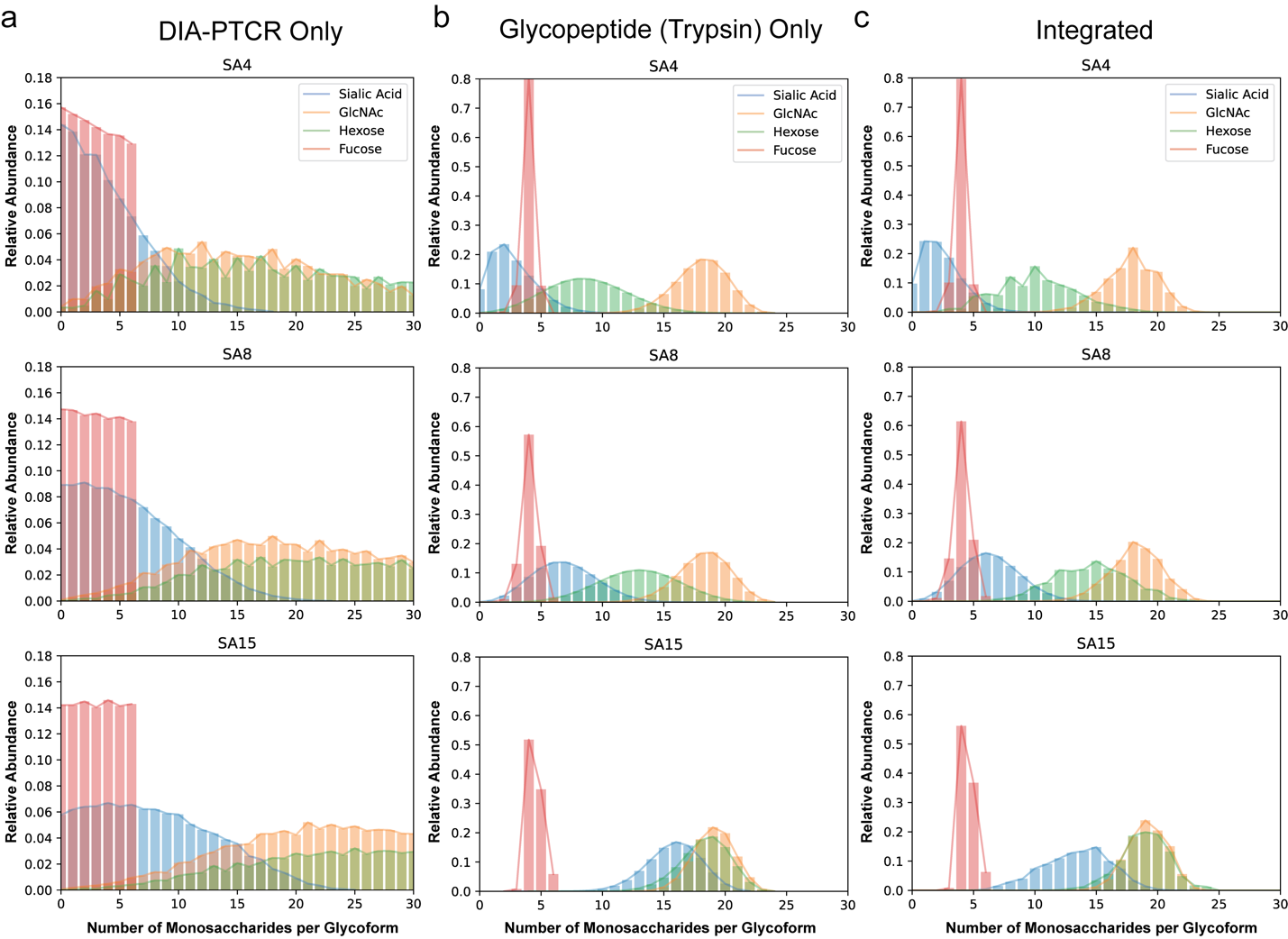

**Supplemental Figure 9.** Number of monosaccharides (i.e., sialic acid, GlcNAc, Hexose and Fucose) per glycoform plotted against their relative abundance in **a**, the DIA-PTCR -only dataset, **b**, the glycopeptide only dataset, **c**, the integrated DIA-PTCR and glycopeptide dataset.

The DIA-PTCR -only plots (**a**) show that without constraining the search space (using glycopeptide data, for example), the detected masses can be assigned to a wide range of glycoforms with varying monosaccharide distributions. First, searching the detected IL22-Fc masses against a database of Chinese Hamster Ovary cell (CHO) glycans (e.g., 180 most common structures) that explain up to 8 glycosylation sites, amounts to >15 x10^9 combinations, which is computationally expensive and highly time-consuming. Second, there are many alternative explanations for the detected glycoforms given their closely related masses. Nonetheless, the DIA-PTCR-only analysis still reveals trends in the sialic acid, fucose and hexose content of the samples that are consistent with the known levels of sialic acid enrichment (Supp. Fig. 8) and orthogonal glycomic analysis (Supp. Fig. 11).

The Trypsin-digested glycopeptide distributions (**b**) show the purely stochastic combinations of all detected monosaccharides from the glycopeptide analysis. Glycopeptide datasets can thus be used to limit the assignments to the most likely glycoforms.

Underpinning this workflow, the integrated DIA-PTCR and glycopeptide data plots (**c**), are determined from the sum of all DIA-PTCR peaks with that assignment, proportionally filtered using glycopeptide data. With the constrained search space, trends in sialic acid and hexose content are further clarified, providing a more precise window into the monosaccharide composition of the samples as a function of sialic acid enrichment.

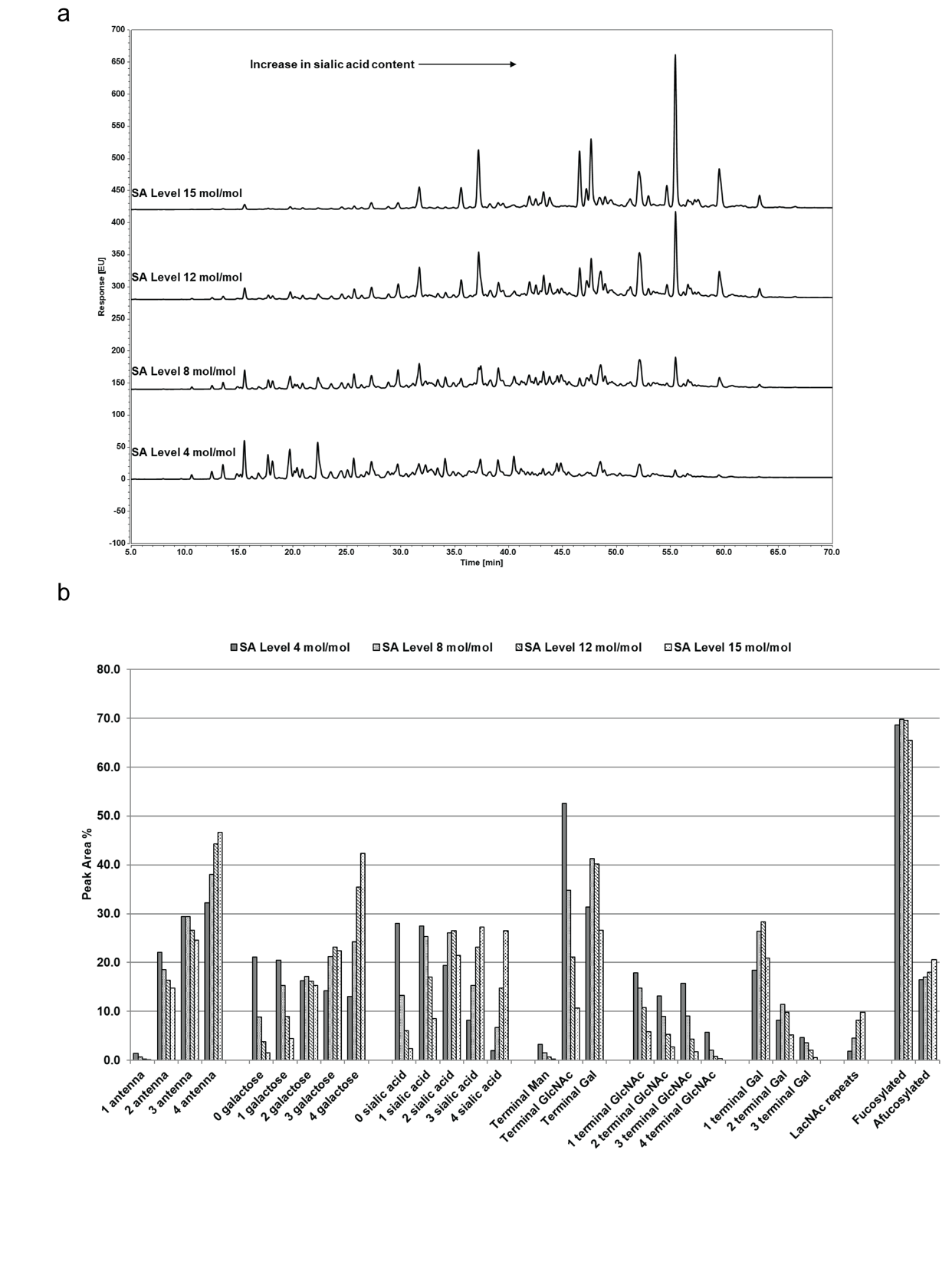

**Supplemental Figure 10. a**, Released glycans and separation of IL22-Fc SA variants by 2AA-labeled N-glycan hydrophilic interaction liquid chromatography (2AA-HILIC). The HILIC method was used to determine the relative distribution of N-linked glycans in IL-22Fc. N-linked glycans were enzymatically released using PNGase F, derivatized with 2-aminobenzoic acid (2-AA), and analyzed by HILIC ultra-high performance liquid chromatography (UHPLC) with fluorescence detection. The chromatograms of the N-linked glycans observed in the sialic acid variants are shown in the above figure. HILIC is a variation of normal phase chromatography in which retention is due to the size and polarity of glycans. The early eluting peaks represent smaller, neutral glycans and the late eluting peaks represent larger, sialylated glycans. As sialic acid content increases, an increase in number of peaks and peak areas for the larger, sialyated glycans is observed along with a corresponding decrease in number of peaks and peak areas for the smaller, neutral glycans. **b**, Breakdown of glycan attributes for SA variants of IL22-Fc as measured by 2AA-HILIC.

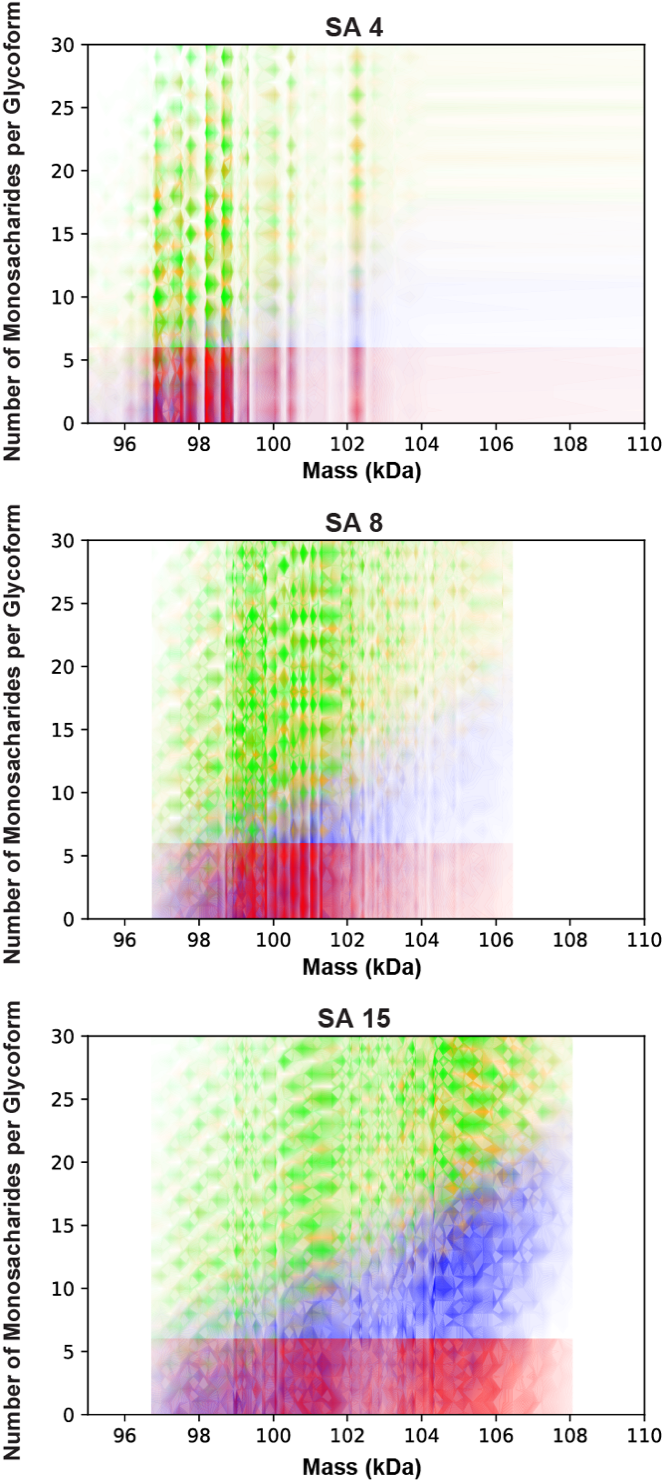

**Supplemental Figure 11.** Number of glycan units/monosaccharides (sialic acid, blue; hexose, green; GlcNAc, yellow; fucose, red) per glycoform plotted against the intact glycoform masses detected by DIA-PTCR for sialic acid (SA) 4, 8 and 15 mol/mol-enriched samples, without constraining the search space using glycopeptide data.

It is noteworthy that trends in sialic acid and hexose content are still apparent in this intact MS-only dataset, which shows that higher mass glycoforms contain higher amounts of sialic acid and hexose with increasing SA enrichment.

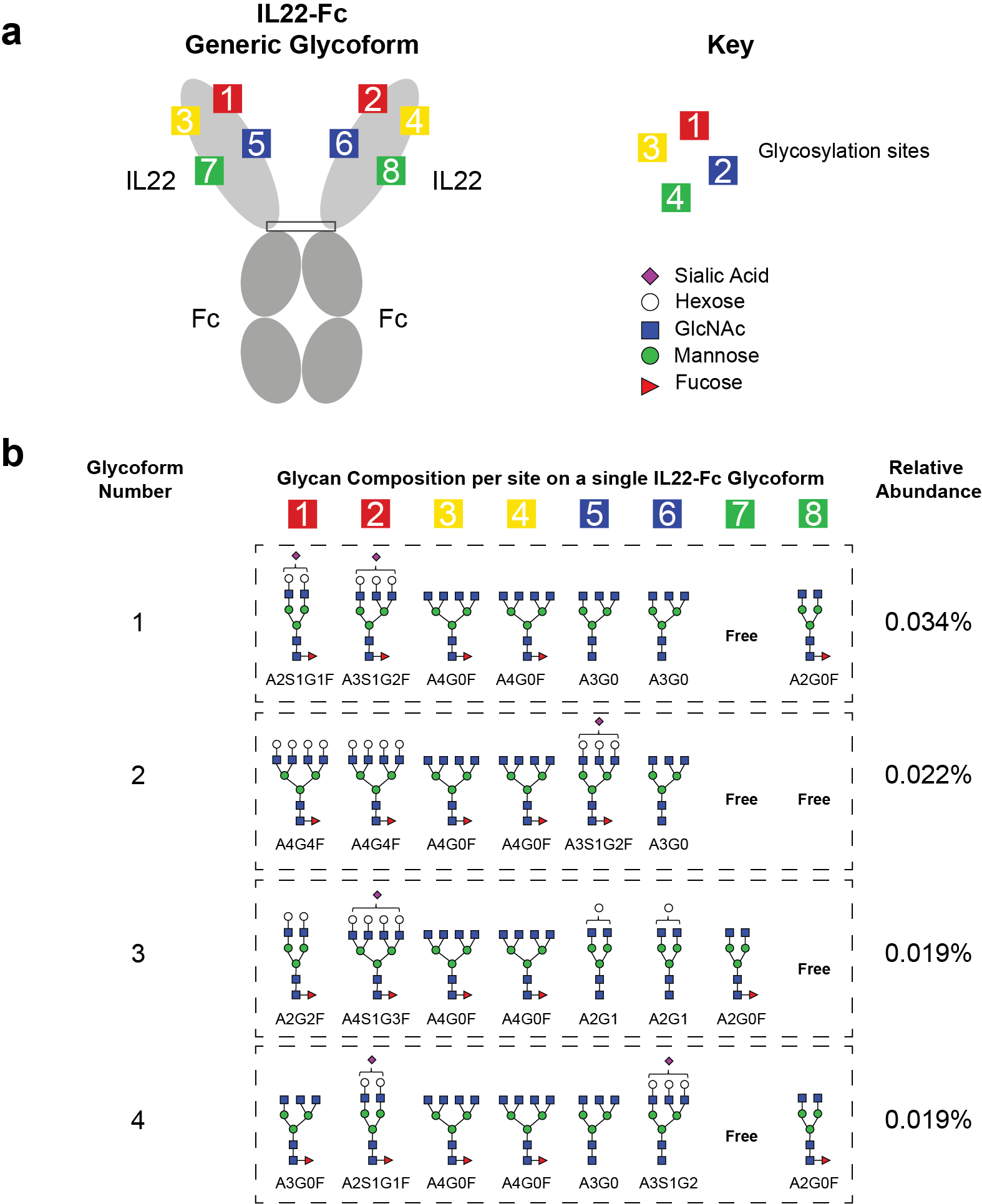

**Supplemental Figure 12. a**, Schematic of IL22-Fc and its 8 glycosylation sites, with a key for the glycan nomenclature. **b**, The four most abundant glycan compositions for each of the 8 possible glycosylation sites on the highest abundance glycoform mass (98,224 Da) detected for IL22-Fc SA4. There are >600,000 other possible glycan compositions – listed in the supplemental table – that explain the same mass peak within 5 Da (the four presented in this figure are the top 4 abundant proteoforms in the list). These findings expose the staggering complexity and heterogeneity of glycoproteins. This is significantly higher than the number of potential monosaccharide annotations because many combinations have the same total monosaccharide composition. Glycoform number 1 in this list corresponds to a single IL22-Fc dimer containing the glycans A2S1G1F, A3S1G2F, 2x A4G0F, 2x A3G0 and A2G0F on seven sites, with the eighth site free / unmodified. For comparison, the highest abundance glycoform composition described above (0.034%) is about nine orders of magnitude more abundant than the lowest abundance glycoform composition for this molecular weight. **Supplemental Spreadsheets 2-4** contain the annotations for detected molecular weights in the all three SA variant conditions.

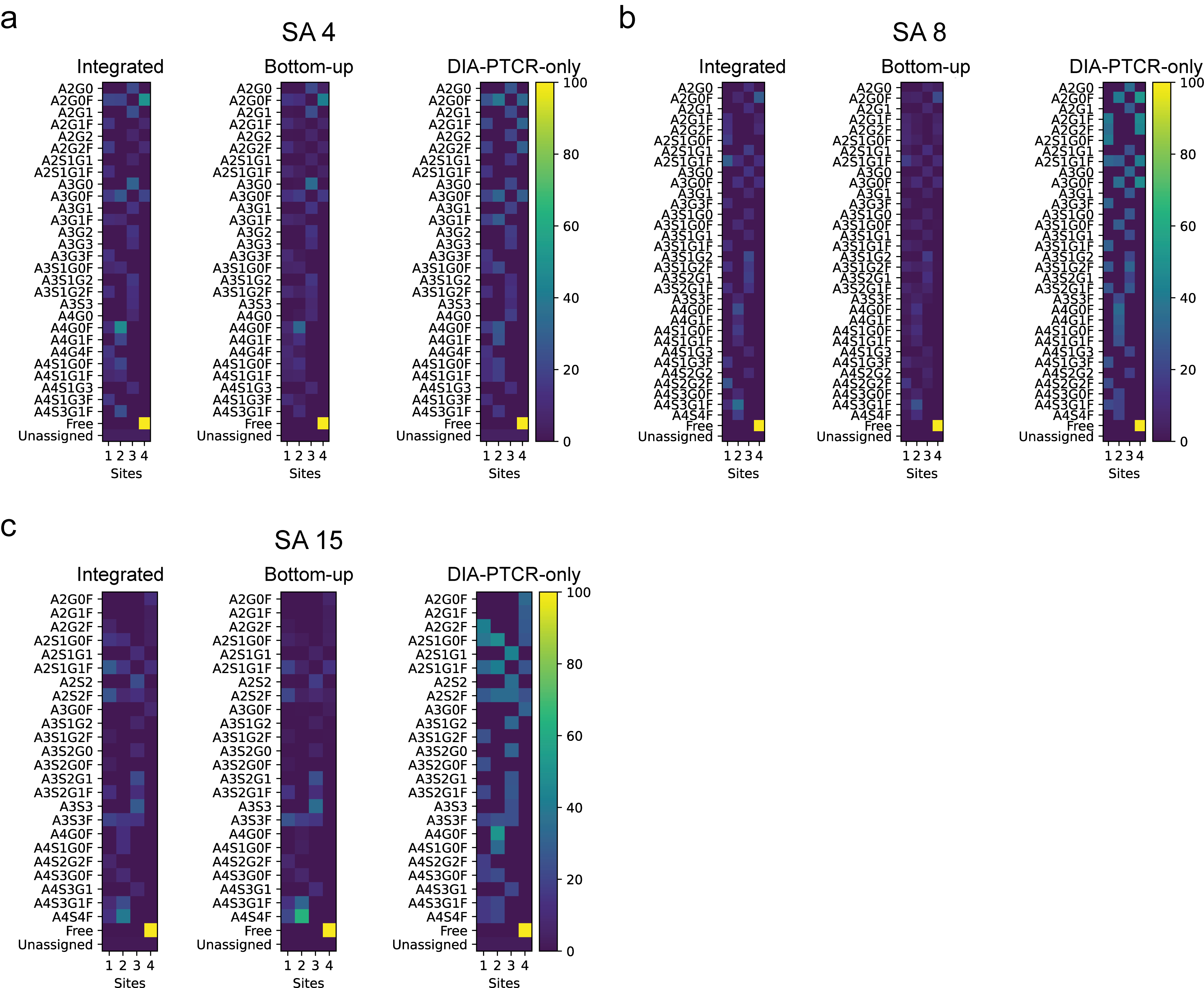

**Supplemental Figure 13.** Glycan barcodes for IL22-Fc SA 4, 8 and 15 (**a,b,c**), showing probability of assignments per site, calculated from the integrated DIA-PTCR and Bottom-up datasets, bottom-up only dataset, and DIA-PTCR-only dataset. There is good agreement between all three barcodes. The DIA-PTCR-only assignments appear to over-represent the probability of site occupancy but are qualitatively comparable to the assignments filtered with glycopeptide-derived probabilities.

To generate these “glycan barcodes”, the probabilities for each potential glycan on each potential site were summed across all peaks, with the relative probability for each potential peak assignment multiplied by the relative peak height from the DIA-PTCR data. Each of these was created from the 8-site glycopeptide data, with a 3% threshold used for the SA4 and SA8 data and a 1% threshold used on the SA15 data. Smaller thresholds were not possible due to memory limitations.

For comparison, and to emphasize the importance of integrating the glycoproteomics and DIA-PTCR data, similar barcodes were made for the DIA-PTCR data alone, which assumed an equal probability for all possible peak assignments that matched within ±5 Da, and from the glycopeptide data alone, without including peak heights from the DIA-PTCR data.

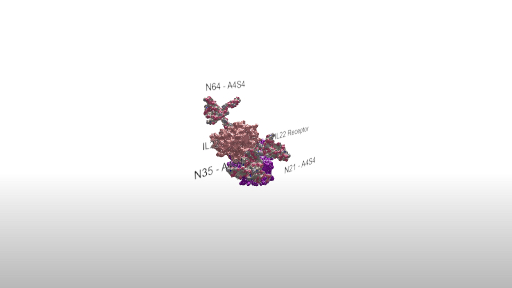

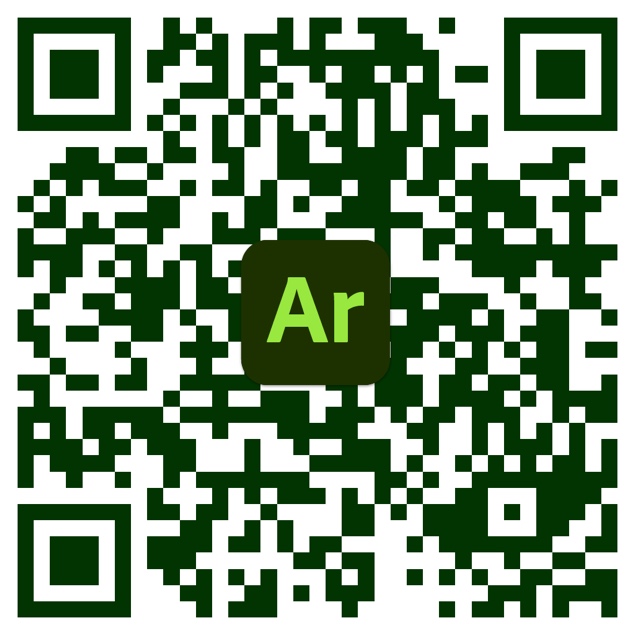

**Supplemental Figure 14.** Augmented reality (AR) structural model of glycosylated IL22 bound to its receptor.

To access the AR experience:

1. Scan the QR code with a smartphone (iPhone, iPad and supported Android devices).
2. Follow prompt to download the Adobe Aero app.
3. Re-scan QR code. Having an Adobe user account is not necessary to access the AR experience.
4. Follow prompt to ‘anchor’ the model to an open space, such as a desk or floor by tapping on the screen.
5. Once anchored, move smartphone around the model to inspect it.
